# Academic criteria for promotion and tenure in faculties of biomedical sciences: a cross-sectional analysis of 146 universities

**DOI:** 10.1101/802850

**Authors:** Danielle B Rice, Hana Raffoul, John PA Ioannidis, David Moher

**Affiliations:** Department of Psychology, McGill University, Montreal, Quebec, Canada; Ottawa Hospital Research Institute, Ontario, Canada; Faculty of Engineering, University of Waterloo, Waterloo, Ontario, Canada; Departments of Medicine, Stanford University, Stanford, California, USA; Health Research and Policy, Stanford University, Stanford, California, USA; Biomedical Data Science, and Statistics, Stanford University, Stanford, California, USA; Meta-Research Innovation Center at Stanford (METRICS), Stanford University, Stanford, California, USA; Centre for Journalology, Clinical Epidemiology Program, Ottawa Hospital Research Institute, Ontario, Canada; School of Epidemiology and Public Health, University of Ottawa, Ottawa, Canada

## Abstract

**Objectives:** To determine the presence of a set of pre-specified traditional and progressive criteria used to assess scientists for promotion and tenure in faculties of biomedical sciences among universities worldwide.

**Design:** Cross-sectional study.

**Setting:** Not applicable.

**Participants:** 170 randomly selected universities from the Leiden Ranking of world universities list were considered.

**Main outcome measures:** Two independent reviewers searched for all guidelines applied when assessing scientists for promotion and tenure for institutions with biomedical faculties. Where faculty-level guidelines were not available, institution-level guidelines were sought. Available documents were reviewed and the presence of 5 traditional (e.g., number of publications) and 7 progressive (e.g., data sharing) criteria was noted in guidelines for assessing assistant professors, associate professors, professors, and the granting of tenure.

**Results:** A total of 146 institutions had faculties of biomedical sciences with 92 having eligible guidelines available to review. Traditional criteria were more commonly reported than progressive criteria (t(82)= 15.1, p= .001). Traditional criteria mentioned peer-reviewed publications, authorship order, journal impact, grant funding, and national or international reputation in 95%, 37%, 28%, 67%, and 48% of the guidelines, respectively. Conversely, among progressive criteria only citations (any mention in 26%) and accommodations for extenuating circumstances (37%) were relatively commonly mentioned; while there was rare mention of alternative metrics for sharing research (2%) and data sharing (1%), and 3 criteria (publishing in open access mediums, registering research, and adhering to reporting guidelines) were not found in any institution reviewed. We observed notable differences across continents on whether guidelines are accessible or not (Australia 100%, North America 97%, Europe 50%, Asia 58%, South America 17%), and more subtle differences on the use of specific criteria.

**Conclusions:** This study demonstrates that the current evaluation of scientists emphasizes traditional criteria as opposed to progressive criteria. This may reinforce research practices that are known to be problematic while insufficiently supporting the conduct of better-quality research and open science. Institutions should consider incentivizing progressive criteria.

**Registration:** Open Science Framework (https://osf.io/26ucp/)

**What is already known on this topic:**

- Academics tailor their research practices based on the evaluation criteria applied within their academic institution.
- Ensuring that biomedical researchers are incentivized by adhering to best practice guidelines for research is essential given the clinical implications of this work.
- While changes to the criteria used to assess professors and confer tenure have been recommended, a systematic assessment of promotion and tenure criteria being applied worldwide has not been conducted.

**What this study adds:**

- Across countries, university guidelines focus on rewarding traditional research criteria (peer-reviewed publications, authorship order, journal impact, grant funding, and national or international reputation).
- The minimum requirements for promotion and tenure criteria are predominantly objective in nature, although several of them are inadequate measures to assess the impact of researchers.
- Developing and evaluating more appropriate, progressive indicators of research may facilitate changes in the evaluation practices for rewarding researchers.

## INTRODUCTION

There are important deficiencies in the quality and transparency of research conducted across disciplines. ^1 2^ Numerous efforts have been made to combat these inadequacies by developing, for example, reporting guidelines (e.g., the CONSORT and PRISMA Statements), registration of studies prior to data collection (e.g., clinicaltrials.gov), and data sharing practices.^3 4^ Despite these strategies, poorly conducted and inadequately reported research remains highly prevalent.^5^ This has important consequences, especially in the field of medicine, as research is heavily relied upon to inform clinical decision-making.

Institutions have the ability to influence large-scale improvements among researchers, as universities hire new faculty, and promote and tenure existing faculty. Universities can provide incentives and rewards (e.g., promotions) for scholarly work that is conducted appropriately, reported transparently, and adheres to best publication practices. A recent survey conducted in the UK found that academics tailor their publication practices to align with their institutional evaluation criteria.^6^ These criteria, however, may include metrics that are known to be problematic for assessing researchers.^7^ Current incentives and rewards may also be misaligned with the needs of society. Reward systems within universities typically incentivize quantity of publications and novelty of findings rather than the reliability, accuracy, reproducibility and transparent reporting of findings.^8^ Inappropriate incentives can inadvertently contribute to research waste,^9^ with billions of dollars invested in non-usable research.^10^ For example, universities that emphasize the quantity of published papers, can increase undeserved authorship, salami slicing, and publication in very low-quality journals (e.g., predatory journals) without peer-review, and contribute to the problems of reproducibility.

Institutional criteria for promotion and tenure decisions can vary and may not be evidence-based.^11^ Some institutions set minimum quantitative “thresholds” for promotion, while others provide qualitative phrasing of criteria that scientists must meet. Recent articles identifying the limitations of the current criteria used to assess scientists for promotion and tenure have been largely conceptual in nature and have limited empirical evidence.^11-14^ In a recent study, evaluation documents were reviewed, and potential limitations were identified.^14^ Further, strategies to encourage implementation and uptake of more responsible indicators for assessing scientists were also discussed. Prior to implementing changes to existing criteria, we must better understand the current standards. Understanding the variability of criteria and thresholds for promotion and tenure applied across institutions requires a systematic empirical assessment. Therefore, we aimed to identify and document a set of pre-specified traditional and progressive criteria used to assess scientists for promotion and tenure within faculties of biomedical sciences among a large number of universities around the world.

## METHODS

The protocol for this study was registered within the Open Science Framework database (https://osf.io/26ucp/) prior to the study’s data collection. The STROBE Statement for cross-sectional studies checklist was used to ensure that methods and findings are clearly reported (see Appendix 1).^15^

### Approach to Selecting University Institutions

The Centre for Science and Technology Studies (CSTS) Leiden Ranking of world universities from 2017 (http://www.leidenranking.com/ranking/2017/list) was used to select institutions for inclusion in the study.^16^ A random convenience sample of 20% of institutions from the Leiden ranking list (170/854) were selected using an online random sampling software.^17^ The CSTS ranking list for the field of “Biomedical and Health Sciences” was selected, which represents the field that publications from universities are assigned to. We planned to include all randomly selected institutions on this list, regardless of the faculties present in each university. The default settings on the CSTS website were used which includes indicator settings of: type of indicator (‘impact’), indicators [‘P, P(top 10%), PP(top 10%)’], which representing the number and proportion of a university’s publications that compared with other publications in the same field and in the same year, are among the top 10% most frequently cited, ordered by publications, and the calculation of impact indicators using fractional counting option selected. A minimum publication output was set at the default value of 100.

### Searching of Institution Criteria

Searching for institutional criteria involved an iterative process by two reviewers. Each institution website was searched for their criteria and policies used for evaluation, promotion and tenure in the faculty of medicine or biomedical sciences faculty. We first searched if each institution had a relevant biomedical sciences department or faculty (e.g., faculty of medicine, department of science). If a faculty of biomedical sciences was found, we searched for keywords on the faculty websites including “academic performance”, “career mobility”, “criteria”, “evaluation”, “guidelines”, “policy”, and “tenure” and “promotion” to find documents related to promotion and tenure. If there was no faculty related to biomedical sciences, or if promotion and tenure guidelines were not publicly available at the faculty level, we referred to the available institution level guidelines. If publicly available criteria could not be located after searching with these methods, human resource personnel, professors, and academic affairs administrators for the institution were contacted directly on up to two occasions to request access to faculty or institution level criteria. Within some countries, promotions first require meeting criteria set at a state or national level.

After searching for faculty- and institution-level guidelines if these were not available, we also searched for state or national guidelines by applying the same search techniques used for universities. For institution websites that were published in languages other than English or French, the promotion and tenure information available on the institution website was searched by an individual who was fluent in the relevant language to facilitate data collection and emails were sent in the language of the institution website. Twelve translators searched university, regional, and national websites for documents to facilitate data extraction. These individuals also translated an email to send to institution representatives when guidelines could not be found online.

### Approach to Selecting List of Criteria

Twelve criteria related to promotion and tenure were purposively selected to enable a comparison between traditional (e.g., quantity of research) and progressive (e.g., reproducibility of research) criteria used to assess scientists for promotion and tenure (see Box 1). Although the characterization of traditional and progressive criteria was ultimately subjective, we based our decisions on evidence and policy initiatives from multiple jurisdictions ^12 14^. We used an iterative process to select the characteristics. An early version of the criteria included 10 items, however, after a set of 5 institutions were pilot tested, two additional items were added, and revisions were made to the wording of select items. The final set of criteria included 5 traditional criteria (i.e., peer-reviewed publications, authorship order, journal impact, grant funding, national or international reputation) and 7 progressive criteria (i.e. citations, data sharing, publishing in open access mediums, registration of research, adherence to reporting guidelines, alternative ways for sharing research, accommodations for extenuating circumstances).

### Data Collection

For each eligible institution, we extracted the following information: university name, faculty name, country, and human development index rating of country.^16^ Faculty of biomedical sciences or institution guidelines used for the evaluation of professors were reviewed, where available, to determine if each of the 12 items from our list of criteria for faculty promotion and tenure were present. We recorded the relevant mentions for each criterion, regardless of how exactly the criterion was considered or operationalized. We did not intend to arbitrate if the proposed version of the criterion was appropriate and technically sophisticated; however, we collected information about if guidelines applied thresholds for each criterion.

We extracted this information for each level of promotion criteria published for universities, including promotions to assistant professor, associate professor, full professor, and the granting of tenure, as well as whether a criterion was mentioned for at least one of these levels. Where institutions applied different labels to ranks/positions (e.g., researcher level A), we sought documentation for the appropriate equivalent categorization of the promotion levels. This information was extracted for tenure-track positions rather than non-tenure track, or clinical professor positions, where available. The level of the promotion criteria available (i.e. faculty level criteria, departmental level criteria), the year that the criterion was published, the associated URL of the criteria, and the date that the website was searched were also extracted. Two reviewers (DBR, HR) independently extracted all data and results were compared for consistency. Where consensus was not achieved between reviewers after discussion, a third team member (DM) was consulted to address discrepancies. For guidelines published in languages other than English or French, translation of the relevant documents was conducted by one individual and verified by a second reviewer (DBR) using Google Translate. A Hungarian translator was not available for one guideline. For this guideline, one reviewer (DBR) used Google translate to conduct data extraction. Data collection was performed using a standardized electronic data collection form in Distiller Systematic Reviewer (Evidence Partners, Ottawa, Canada).

### Approach to Synthesis

Institution characteristics and promotion and tenure criteria were summarized in table format to facilitate inspection and discussion of findings. The percentage of criteria that were included in promotion and tenure guidelines were summed and presented in table format and means for traditional and progress criteria were compared through a paired sample t-test after ensuring the data was normally distributed. Categorical variables were presented as percentages and counts, and continuous variables were presented as means and standard deviations (SD). Institutions that had criteria available were compared to institutions that did not have criteria available through independent samples t-test and chi-square tests.

Exploratory analyses were conducted for the full professor position as this had the most data available. Two multiple linear regressions were conducted to assess the associations between institution characteristics (level of criteria, continent, human development index, and Leiden ranking) and the number of criteria present for traditional and progressive items for guidelines for professors as the majority of institutions had guidelines present for this promotion level. Logistic regressions were conducted for each criterion present among 10% to 90% of institutions at the rank of full professor. Prior to conducting regression analyses, preliminary tests were performed to confirm that there were no violations of multiple regression assumptions. Assumptions of outliers, normality, linearity, homoscedasticity, and independence of residuals were checked by inspection of the normal probability plot of the regression standardized residual and the scatter plot. Microsoft Excel was used for summing study results, while SPSS Statistics version 21.0 (Chicago, IL) was used for statistical tests. All statistical analyses were two-tailed with *p*<.005 significance level to adhere to recent recommendations for a lowered threshold of statistical significance.^14 18^

Five institutions (University of Valencia, Case Western Reserve University, University of Nebraska – Lincoln, University of Florence, and University of Maryland – Baltimore) were randomly selected to request feedback on the documents used for data extraction and the accuracy of data extraction. An email was sent to the deans of universities in addition to the associate dean of research (or equivalent) in the relevant medical faculty. A confirmation of email receipt was requested, and a reminder email was sent if there was no response after two weeks.

### Patient and Public Involvement

This research did not involve patients or the public.

## RESULTS

### Deviations from Protocol

We refined our inclusion criteria to exclude institutions that did not have a faculty of medicine, biomedical sciences, life sciences, health sciences, or medical sciences in order to focus on institutions that have a department directly related to studying and subsequently disseminating biomedical research.

### Institution Characteristics

Case Western Reserve University and the University of Florence responded to our request for reviewing the accuracy of our data extraction. The university representatives considered our data an accurate reflection of their policies with one caveat from the University of Florence that the criteria are modified from “time to time by the commission”. Of the 170 institutions reviewed, 146 had faculties of biomedical sciences. We were able to access a total of 92 (63%) institutions’ guidelines for promotion and tenure (see Appendix 2 and Figure 1). For the other institutions we could not find or access such guidelines either online or after communication (see Appendix 3). Four institutions responded to email inquiries and noted that they do not have such guidelines (Karolinska Institute, Heidelberg University, Freie Universitãt Berlin, and University of Tartu). Of institutions with available guidelines, 39 (42%) were specific to faculties of biomedical sciences.

The universities that we could evaluate had mostly a very high development index rating (n=68, 74%) and they were almost equally split between Europe (n=27, 29%), Asia (n=29, 32%), and North America (n=28, 30%). Guidelines referred to were last updated between 1993 and 2018 (median = 2016, interquartile range [IQR] 2011 to 2017). Based on Leiden ranking of world universities, institution rankings ranged from 6 to 842 (median = 345, IQR 157 to 549) (see Table 1). Of the 92 guidelines reviewed, the evaluation for promotion to assistant professor (n=49, 53%), associate professor (n=79, 86%), full professor (n=83, 90%), and tenure (n=26, 28%) were present, with most (n=82, 89%) institutions having guidelines available for more than one level of promotion. There were no statistically significant differences between institutions that did versus did not have criteria available for institution rankings (t(144) = -1.48, *p* = .142) or human development index (x^2^(2, *n* = 93, *p* = .918). We observed notable differences across continents on whether guidelines were accessible or not (Australia 100%, North America 97%, Europe 50%, Asia 58%, South America 17%) (x^2^(5, *n* = 93, *p* = .001) (see Table 1).

The traditional incentives that were present most often were peer-reviewed publications (any mention: n=87, 95%; assistant professor: n= 39, 80%; associate professor: n=76, 96%; professor: n=79, 95%; tenure: n=22, 85%) and grant funding (any mention: n=62, 67%; assistant professor: n=26, 53%; associate professor n=50, 63%; professor: n=56, 67%; tenure: n=15, 58%) with less frequent use of authorship order (any mention: 34, 37%), journal impact (any mention: n=26, 28%), and national or international reputation (any mention: n=44, 48%). The exact mentions and how they were supposed to be operationalized varied across guidelines and only a minority were quantitative. Thirty-two institutions (35%) had at least one mention to a specific number of peer-reviewed publications. The requirements for publications was heterogeneous between institutions with some, for example requiring a specific number per year, or over the past 10 years (e.g., as few as 1 total publication required with as many as 53 publications required in the past 10 years). Institutions that required fewer publications often reported that publishing in journals with higher impact factors was necessary (i.e., one publication in a journal with an impact factor of at least 10, or two publications in journals with impact factors of at least 5). Fourteen institutions (15%) had at least one mention to a specific amount of money for funding [range 300,000 Rem Min Bi (41,766 USD) to 3,000,000 Rem Min Bi (417,660 USD]. For authorship order, 24 (26%) institutions encouraged first author publications, 20 (22%) encouraged last or corresponding author publications, although many of these institutions also promoted first author publications, and 4 institutions (11%) encouraged sole authored publications. No institutions mentioned that middle author publications or multi-authored papers were favourable. For journal impact, 11 institutions of 26 (42%) mentioned specific numbers for desirable impact factor metrics, but the desirable impact factor thresholds varied enormously across institutions (≥ 3, 4, 5, 9, 10, 11, 20, 30, see Appendix 4). No institutions had any numerical recommendations on assessment of national or international reputation.

For the 7 progressive items, only 3 incentives (43%) were present in at least one institution for every level of promotion guidance. Progressive incentives that were present included, adjustments to expectations when professors go on leave (any mention: n=34, 37%; assistant professor: n=22, 45%; associate professor: n=28, 35%; professor: n=29, 35%; tenure: n=13, 50%), citations of research (any mention: n=24, 26%; assistant professor: n=12, 24%; associate professor: n=23, 29%; professor n=23, 28%; tenure: n=6, 23%), and rarely mention of alternative metrics for sharing research (any mention: n= 3, 3%; assistant professor: n=3, 6%; associate professor: n=3, 4%; professor: n=2, 2%; tenure: n=1, 4%). Data sharing was mentioned only in 1 institution (1%). Mention of publishing in open access outlets, registering research, and adhering to reporting guidelines were absent from all institutions (see Table 2 and Figure 2). Progressive criteria were mostly qualitative. For citations, however, 25% of institutions (6 of 24) that included this item proposed specific numbers (see Appendix 4).

### Characteristics Associated with the Presence of Traditional and Progressive Items for Professors

In tests of multicollinearity, independent variable tolerance values ranged from 0.4 to 0.9, and the variance inflation factors ranged from 1.1 to 2.8 for both traditional and progressive analyses, indicating that multicollinearity was not a major issue.^19^ There was no deviation in the assumption of normality based on the inspection of the normal probability plots of the regressions or evidence of violations of assumptions of outliers, linearity, homoscedasticity, and independence of residuals based on the standardized residual and scatter plot inspections.

#### Regressions for Total Scores

The mean number of traditional items present for promotion criteria used for full professors was 2.7 (SD = 1.2) of 5 criteria. Institutions from Australia (standardized regression coefficient [β] = 0.39, *p* = .004) and North America (β = 0.39, *p* = .025) tended to have a slightly larger number of traditional criteria present in guidelines (Table 3). Australia had an average of 3.5 (SD = 1.0) traditional criteria present while North America had an average of 3.2 (SD = 1.1). For progressive criteria, the mean number of items present was 0.7 (SD = 0.8) of 7 criteria. A significantly greater number of traditional items compared to progressive items were reported among institutions (t(82)= 15.1, p= .001). Institutions located in Australia (β = 0.37, *p* = .006) had modestly more progressive criteria in their guidelines (Table 3). Institutions from Australia had an average of 1.5 (SD=0.5) progressive criteria present in guidelines.

#### Regressions for Individual Items

Six of the 12 items were present in 10% to 90% of promotion and tenure guidelines for professors, including grant funding, authorship order, impact factor, national or international reputation, citations, and adjustments to expectations. Encouraging researchers to have a national or international reputation was significantly more present among institutions from North America (β = 27.44, 95% confidence interval [CI] = 3.26, 231.16, *p* = .002). Australia (β = 52.36, 95% CI = 2.03, 1190.95, *p* = .013) tended to have a slightly larger number of institutions encouraging citations in promotion and tenure guidelines (Table 4).

## DISCUSSION

We found that traditional criteria are relied on in guidelines for assessing faculty members for promotion and tenure among an international sample of 92 institutions with faculties of biomedicine or health sciences. Almost all institutions incentivized the presence of peer-reviewed publications, many of which also required a minimum number of papers published per year. Conversely, only about a third of institutions discussed citations and none referenced publishing in open access mediums, registering research, and adhering to reporting guidelines for transparently presenting research was absent from all examined documents.

There was substantial variability across continents on whether any guidelines were available at all. Given the substantial rate of non-response, we cannot exclude the possibility that such documents exist but could not be retrieved. For some universities, the criteria and related guidelines are not set at the level of the medical faculty or even the whole university, but by a higher state authority (e.g. the ministry of education in Greece) for all universities. Availability of criteria is probably helpful for transparency; however, the availability of guideline documents does not mean that these are also faithfully adhered to.

An important barrier to the implementation of progressive criteria relates to the difficulty of selecting and integrating more appropriate measures,^14^ which was described in several promotion documents reviewed. Institutions have noted the imperfections of traditional criteria being applied for assessing professors such as the impact factor, but reported that few alternatives exist.^20 21^ Integrating progressive criteria to incentivize scientists requires evidence on the accuracy and the validity of progressive indicators ^14^; and such indicators are starting to emerge.

As progressive metrics are available, implementing their use more widely, for example through Declaration on Research Assessment (DORA)’s advisory board, may be one avenue to aid in the dissemination of more appropriate tools for assessing scientists.

Institutions relying on traditional metrics, such as number of publications and associated journal impact factors may misinterpret what these metrics mean.^21^ Beyond evidence, there are other reasons to consider alternative criteria. They may better align with a university’s mission. Similarly, some criteria, such as data sharing, has a high research integrity value; patients support sharing of their data ^22^ and it facilitates assessments of reproducibility. To facilitate data sharing it is likely that the FAIR (Findability, Accessibility, Interoperability and Reusability) principles will need to be in place.^9^ An additional difficulty to including progressive criteria in evaluations is the need for resources to support this change. The institution reviewed in our sample that had the greatest number of progressive criteria, Ghent University, described having invested in an online system to help assess some progressive metrics of their researchers.^20^ Decreasing the barriers to using progressive metrics in evaluations will be necessary for systematic changes to occur. There are limitations that should be considered when interpreting our study results. Despite searching websites and contacting institutions, not all institutions use prespecified criteria for assessing promotion and in some instances, documents were not found. A second limitation is that incentives for professors can occur through other pathways, e.g. financial bonuses, which may not be publicly available or included in the documents reviewed. Obtaining a more complete understanding of the criteria used for providing financial and reputational incentives in medical faculties may include reviewing internal documentation regarding bonuses and awards or recognitions in addition to formal promotions. Furthermore, medical faculties often take into account clinical work and teaching which we did not include.

Finally, we should acknowledge that both for traditional and for progressive criteria, the exact way they are proposed and operationalized can make a difference on whether they can have a positive or negative impact on research quality. With a plethora of metrics being developed for progressive criteria, some of them may be much better than others. Integrating appropriately framed criteria that incentivize excellence and high quality could result in improvements in medical research and evidence-based medicine. Systematic changes require collaborative efforts and creativity in order to address barriers to developing and adopting the best metrics. Considering the benefits of creating sustainable changes to the incentives that drive poor medical research internationally, however, would be a turning point in facilitating the transparency, openness, and reproducibility in research practices.

### Box 1.

**Criteria of Interest for Promotion and Tenure.**

#### Traditional Incentives

1. Is any quantitative or qualitative mention made about publications required? If quantitative, please specify the requirement.
2. Is any quantitative or qualitative mention made about the specific authorship order in publications? If so, please specify order (e.g. first, senior, single) required.
3. Is any mention made of journal impact factors? If quantitative, what are the minimum thresholds?
4. Is any mention made of grant funding? If quantitative, what are the minimum thresholds (i.e., amount of funding and/or number of grants as principal investigator)?
5. Is any mention made requiring that research is recognized at a national or international level? If so, please specify the requirement.

#### Progressive Incentives

6. Is any mention made of citations? If quantitative, what are the thresholds of minimum requirement? Are specific citation databases mentioned?
7. Is any mention made of data sharing? If quantitative, what are the minimum thresholds (e.g., percentage of data that is to be made available)?
8. Is any mention made of publishing in open access mediums? If quantitative, what are the minimum thresholds (e.g., percentage of studies to be published in open access journals)?
9. Is any mention made of registration (including preregistration challenge) of studies? If yes, are there thresholds of minimum requirement (e.g., percentage of studies that are to be registered).
10. Is any mention made of adherence to reporting guidelines for publications? If so, are specific guidelines mentioned?
11. Is any mention made of alternative metrics for sharing research (e.g., social media and print media)? If so, are specific metrics mentioned?
12. Is any mention made of accommodations or adjustments to expectations due to extenuating circumstance? If so, please specify the description of accommodations (e.g., an extra year to defer tenure consideration) and the type of eligible circumstances (e.g., parental leave, medical leave)?

## ACKNOWLEDGEMENTS

We would like to thank Dr. Juan Pablo Alperin for providing us with access to the database which included documents for review from North American Universities. We would like to thank Becky Skidmore for her recommended search terms and Nik Chander for his help with table formatting. We would also like to thank each of the individuals that searched websites and translated documents including: Xiaoqin Wang, Andrea Carboni Jiménez, Michal Dedys, Fatemeh Yazdi, Philipp-Clemens Nowotny, Song Xiaoyang, Jin Yan, Elena Tarasova, Kednapa Thavorn, Chen He, Alexander Tsertsvadze, and Francesca Ruggiero. Finally, we would like to acknowledge each of the institutions that responded to our requests for promotion and tenure documents.

## ETHICAL APPROVAL

As this study did not involve any human data, ethics approval was not required.

## TRANSPARENCY DECLARATION

The manuscript’s guarantor affirms that the manuscript is an honest, accurate, and transparent account of the study being reported; that no important aspects of the study have been omitted; and that any discrepancies from the study as planned (and, if relevant, registered) have been explained.

## DISSEMINATION DECLARATION

Not application. This work did not involve study participants.

## DATA SHARING

All data associated with this study is posted on the open science framework (https://osf.io/26ucp/). The study protocol, data extraction forms, and data are also available at this link.

## AUTHOR CONTRIBUTIONS

DM and JPAI conceived of the study. DBR and DM wrote the study protocol and the initial draft of the article. DM, DBR and HR agreed on the criteria applied for promotion and tenure after pilot testing a set of university criteria. DBR and HR found institution guidelines and extracted information regarding promotion criteria. All authors were involved in subsequent protocol revisions. DBR conducted the statistical analyses. All authors provided critical feedback and agreed to the final version of the paper.

