## Appendix 1 for "Academic criteria for promotion and tenure in faculties of biomedical sciences: a cross-sectional analysis of 146 universities"

**Appendix 1. Adherence to STROBE Guideline.**

|  | **Item No** | **Recommendation** | **Page Number** |
| --- | --- | --- | --- |
| **Title and abstract** | 1 | (*a*) Indicate the study’s design with a commonly used term in the title or the abstract | 1 |
|  |  | (*b*) Provide in the abstract an informative and balanced summary of what was done and what was found | 2-3 |
| **Introduction** |  |  |  |
| Background/rationale | 2 | Explain the scientific background and rationale for the investigation being reported | 5-6 |
| Objectives | 3 | State specific objectives, including any prespecified hypotheses | 6 |
| **Methods** |  |  |  |
| Study design | 4 | Present key elements of study design early in the paper | 6 |
| Setting | 5 | Describe the setting, locations, and relevant dates, including periods of recruitment, exposure, follow-up, and data collection | N/A |
| Participants | 6 | *Cross-sectional study*—Give the eligibility criteria, and the sources and methods of selection of participants | 6-8 |
| Variables | 7 | Clearly define all outcomes, exposures, predictors, potential confounders, and effect modifiers. Give diagnostic criteria, if applicable | 9-10 |
| Data sources/ measurement | 8* | For each variable of interest, give sources of data and details of methods of assessment (measurement). Describe comparability of assessment methods if there is more than one group | 9-10, Appendix 2 |
| Bias | 9 | Describe any efforts to address potential sources of bias | 11 |
| Study size | 10 | Explain how the study size was arrived at | 6 |
| Quantitative variables | 11 | Explain how quantitative variables were handled in the analyses. If applicable, describe which groupings were chosen and why | 10-11 |
| Statistical methods | 12 | (*a*) Describe all statistical methods, including those used to control for confounding | 10-11 |
|  |  | (*b*) Describe any methods used to examine subgroups and interactions | N/A |
|  |  | (*c*) Explain how missing data were addressed | N/A |
|  |  | *Cross-sectional study*—If applicable, describe analytical methods taking account of sampling strategy | N/A |
|  |  | (*e*) Describe any sensitivity analyses | N/A |

| **Results** |  |  |  |
| --- | --- | --- | --- |
| Participants | 13* | (a) Report numbers of individuals at each stage of study—eg numbers potentially eligible, examined for eligibility, confirmed eligible, included in the study, completing follow-up, and analysed | 12, Figure 1 |
|  |  | (b) Give reasons for non-participation at each stage | Appendix 3 |
| Descriptive data | 14* | (a) Give characteristics of study participants (eg demographic, clinical, social) and information on exposures and potential confounders | 12, Table 1 |
|  |  | (b) Indicate number of participants with missing data for each variable of interest | 11-14 |
| Outcome data | 15* | *Cross-sectional study—*Report numbers of outcome events or summary measures | 11-14 |
| Main results | 16 | (*a*) Give unadjusted estimates and, if applicable, confounder-adjusted estimates and their precision (eg, 95% confidence interval). Make clear which confounders were adjusted for and why they were included | 10-11 |
|  |  | (*b*) Report category boundaries when continuous variables were categorized | N/A |
|  |  | (*c*) If relevant, consider translating estimates of relative risk into absolute risk for a meaningful time period | N/A |
| Other analyses | 17 | Report other analyses done—eg analyses of subgroups and interactions, and sensitivity analyses | 10-11 |
| **Discussion** |  |  |  |
| Key results | 18 | Summarise key results with reference to study objectives | 15-16 |
| Limitations | 19 | Discuss limitations of the study, taking into account sources of potential bias or imprecision. Discuss both direction and magnitude of any potential bias | 17 |
| Interpretation | 20 | Give a cautious overall interpretation of results considering objectives, limitations, multiplicity of analyses, results from similar studies, and other relevant evidence | 15-18 |
| Generalisability | 21 | Discuss the generalisability (external validity) of the study results | 16 |
| **Other information** |  |  |  |
| Funding | 22 | Give the source of funding and the role of the funders for the present study and, if applicable, for the original study on which the present article is based | 23-24 |
