## Appendix 2 for "Academic criteria for promotion and tenure in faculties of biomedical sciences: a cross-sectional analysis of 146 universities"

| **University** | **Level of Criteria Assessed** | **Leiden Ranking of University** | **Most Relevant Biomedical Faculty** |
| --- | --- | --- | --- |
| Bar-Ilan University | Faculty Criteria | 495 | Faculty of Medicine |
| Brigham Young University | Institution Criteria | 565 | Department of Health Sciences |
| Cardiff University | Institution Criteria | 213 | School of Medicine |
| Case Western Reserve University | Faculty Criteria | 62 | School of Medicine |
| Central China Normal University | Institution Criteria | 790 | Life Sciences |
| Central South University | Faculty Criteria | 84 | School of Medicine |
| China Medical University | Faculty Criteria | 96 | Medical University |
| China Pharmaceutical University | Faculty Criteria | 298 | Basic Medicine and Clinical Pharmacy |
| Colorado State University | Faculty Criteria | 400 | School of Medicine |
| Dankook University | Institution Criteria | 575 | College of Medicine |
| Duke University | Faculty Criteria | 15 | School of Medicine |
| Ghent University | Institution Criteria | 94 | Faculty of Medicine and Health Sciences |
| Guangxi Medical University | Faculty Criteria | 328 | Medical University |
| Guangzhou Medical University | Faculty Criteria | 354 | Medical University |
| Gunma University | Faculty Criteria | 440 | Faculty of Medicine |
| Harbin Medical University | Institution Criteria | 152 | Medical University |
| Johannes Gutenberg University Mainz | Faculty Criteria | 232 | University Medicine |
| Kansas State University | Faculty Criteria | 580 | Department of Diagnostic Medicine/Pathobiology |
| Katholieke Universiteit Leuven | Institution Criteria | 56 | Faculty of Medicine |
| Korea Advanced Institute of Science and Technology | Faculty Criteria | 547 | Graduate School of Medical Science and Engineering |
| Kyung Hee University | Institution Criteria | 183 | College of Medicine |
| Lomonosov Moscow State University | Institution Criteria | 533 | Faculty of Fundamental Medicine |
| Murdoch University | Institution Criteria | 761 | School of Veterinary and Life Sciences |
| Nagasaki University | Faculty Criteria | 349 | School of Medicine |
| Nanchang University | Faculty Criteria | 420 | Medical College |
| Nanjing Medical University | Institution Criteria | 72 | Medical University |
| National Autonomous University of Mexico - UNAM | Institution Criteria | 260 | School of Medicine |
| National University of Cordoba | Faculty Criteria | 639 | School of Medicine |
| North Dakota State University | Faculty Criteria | 695 | School of Medicine & Health Sciences |
| Northeast Normal University | Institution Criteria | 767 | School of Life Science |
| Norwegian University of Science and Technology | Institution Criteria | 303 | Faculty of Medicine and Health Sciences |
| Ohio University | Faculty Criteria | 694 | Biomedical Sciences |
| Old Dominion University | Institution Criteria | 728 | College of Health Sciences |
| Philipps-Universität Marburg | Institution Criteria | 348 | Department of Medicine |
| Prince Songkla University | Institution Criteria | 624 | Faculty of Medicine |
| Queen Mary University of London | Institution Criteria | 240 | Barts and the London School of Medicine and Dentistry |
| Seoul National University | Institution Criteria | 14 | College of Medicine |
| Shahid Beheshti University of Medical Sciences | Institution Criteria | 317 | School of Medicine |
| Shenyang Pharmaceutical University | Institution Criteria | 479 | Medicinal Chemistry |
| Shiraz University of Medical Science | Institution Criteria | 423 | School of Medicine |
| South China Agricultural University | Institution Criteria | 629 | College of Life Sciences |
| South China University of Technology | Institution Criteria | 630 | School of Medicine |
| St Louis University | Faculty Criteria | 385 | School of Medicine |
| Sun Yat-sen University | Faculty Criteria | 26 | Medical Sciences |
| Tulane University | Faculty Criteria | 308 | School of Medicine |
| Uniformed Services University of the Health Sciences | Institution Criteria | 421 | School of Medicine |
| Università degli Studi di Genova | Institution Criteria | 253 | School of Medical and Pharmaceutical Sciences |
| Université Catholique de Louvain | Institution Criteria | 257 | Faculty of Medicine and Dentistry |
| University College Dublin | Institution Criteria | 252 | School of Medicine |
| University of Akron | Faculty Criteria | 764 | Medicine |
| University of Belgrade | Institution Criteria | 214 | School of Medicine |
| University of Campinas | Institution Criteria | 186 | School of Medical Sciences |
| University of Chile | Institution Criteria | 415 | Faculty of Medicine |
| University of Colorado, Denver | Faculty Criteria | 51 | School of Medicine |
| University of East Anglia | Institution Criteria | 504 | Medical School |
| University of Florence | Institution Criteria | 177 | Faculty of Medicine and Dentistry |
| University of Freiburg | Institution Criteria | 142 | Faculty of Medicine |
| University of Kansas | Faculty Criteria | 181 | School of Medicine |
| University of Maryland, Baltimore | Faculty Criteria | 98 | School of Medicine |
| University of Melbourne | Institution Criteria | 21 | Medicine School |
| University of Minnesota - Twin Cities | Faculty Criteria | 32 | Department of Medicine |
| University of Modena and Reggio Emilia | Institution Criteria | 397 | Faculty of Medicine and Surgery |
| University of Montpellier | Institution Criteria | 321 | Faculty of Medicine |
| University of Navarra | Institution Criteria | 433 | School of Medicine |
| University of Nevada, Las Vegas | Faculty Criteria | 686 | School of Medicine |
| University of Nevada, Reno | Faculty Criteria | 657 | School of Medicine |
| University of New South Wales | Institution Criteria | 87 | Medicine |
| University of North Texas | Faculty Criteria | 567 | College of Osteopathic Medicine |
| University of Oklahoma | Faculty Criteria | 234 | College of Medicine |
| University of Oslo | Institution Criteria | 73 | Faculty of Medicine |
| University of Pittsburgh | Faculty Criteria | 6 | School of Medicine |
| University of Pretoria | Faculty Criteria | 549 | School of Medicine |
| University of Queensland | Institution Criteria | 50 | Faculty of Medicine |
| University of Saskatchewan | Faculty Criteria | 345 | College of Medicine |
| University of South Australia | Institution Criteria | 437 | School of Pharmacy and Medical Sciences |
| University of St Andrews | Institution Criteria | 689 | School of Medicine |
| University of Strathclyde | Institution Criteria | 619 | Institute of Pharmacy and Biomedical Sciences |
| University of Sussex | Institution Criteria | 555 | Medical School |
| University of Szeged | Institution Criteria | 470 | Faculty of Medicine |
| University of Tampere | Faculty Criteria | 386 | Faculty of Medicine and Life Sciences |
| University of Tennessee, Knoxville | Institution Criteria | 187 | Graduate School of Medicine |
| University of Toledo | Faculty Criteria | 487 | College of Medicine and Life Sciences |
| University of Utah | Faculty Criteria | 80 | School of Medicine |
| University of Valencia | Faculty Criteria | 278 | Faculty of Medicine and Odontology |
| University of Western Australia | Institution Criteria | 178 | Medical School |
| University of Windsor | Institution Criteria | 784 | Schulich School of Medicine and Dentistry |
| Utrecht University | Institution Criteria | 37 | Faculty of Medicine |
| Wuhan University | Faculty Criteria | 138 | Faculty of Medical Sciences |
| Xi'an Jiaotong University | Faculty Criteria | 157 | School of Basic Medical Sciences |
| Xidian University | Institution Criteria | 842 | School of Life Sciences and Technology |
| Yamagata University | Institution Criteria | 634 | Faculty of Medicine |
| Yeungnam University | Institution Criteria | 476 | College of Medicine |
