## Appendix 3 for "Academic criteria for promotion and tenure in faculties of biomedical sciences: a cross-sectional analysis of 146 universities"

**Appendix 4. Categorization of Institutions Not Included in Analyses**

| **Emailed Institution Twice with No Response** | 1. Akdeniz University* 2. Budapest University of Technology and Economics* 3. Chungnam National University 4. COMSATS Institute of Information Technology 5. Ehime University 6. Ernst-Moritz-Arndt University of Greifswald 7. Federal University of São Carlos - UFSCar 8. Fourth Military Medical University 9. Graz University of Technology* 10. Jiangnan University 11. Jilin University 12. Kindai University 13. Kobe University 14. Lille 1 University of Science and Technology 15. Medical University of Graz 16. Metropolitan Autonomous University 17. National Technical University of Athens* 18. National Tsing Hua University 19. National Yang Ming University 20. Nicholas Copernicus University of Torun 21. Osaka University 22. Peking Union Medical College 23. Saarland University 24. Shanghai Jiao Tong University 25. Tokyo Medical and Dental University 26. Tokyo University of Agriculture and Technology 27. Universidad Nacional de La Plata* 28. Universidade Federal do Paraná 29. Universidade Federal do Rio de Janeiro 30. Université Paul Sabatier 31. Universiti Teknologi Malaysia 32. University of Hyderabad 33. University of La Laguna 34. University of Minho 35. University of Paris XII - Paris-Est Créteil Val de Marne 36. University of Silesia 37. University of Southampton 38. University of Thessaly 39. University of Tokyo 40. University of Toyama 41. University of Valladolid 42. Waseda University 43. Wroclaw University of Technology |
| --- | --- |
| **Institution Reported No Promotion or Tenure Criteria is Applied** | 1. Freie Universität Berlin 2. Heidelberg University 3. Karolinska Institute 4. University of Tartu |
| **Institution not Willing to Share Documents** | 1. City University London 2. Erasmus University Rotterdam* |
| **Institution Provided Documents for Hiring Only** | 1. University of São Paulo 2. Tohoku University 3. University of Bonn 4. University of Paris VII - Paris Diderot 5. University of Santiago de Compostela 6. Aalborg University 7. Gifu University 8. Yamaguchi University |
| **Guidelines Only Available for Positions Not Included in Analyses** | 1. University of Münster 2. Julius Maximilian University of Würzburg 3. University of Girona |
| **Documents Available but No Relevant Faculty** | 1. Anna University* 2. Beihang University* 3. Carnegie Mellon University* 4. Delft University of Technology* 5. Eindhoven University of Technology* 6. George Mason University* 7. Georgia Institute of Technology* 8. Hefei University of Technology* 9. Iowa State University* 10. Nanjiang Agricultural University* 11. Roma Tre University* 12. Sichuan Agricultural University* 13. Technische Universität Darmstadt* 14. University of Bath* 15. University of Nebraska, Lincoln* 16. University of Salento* 17. University of Stuttgart* 18. Vienna University of Technology* |

***Note***: * = institutions did not have a relevant biomedical faculty
