## Appendix 4 for "Academic criteria for promotion and tenure in faculties of biomedical sciences: a cross-sectional analysis of 146 universities"

**Appendix 4. Thresholds for Criteria**

| **Criteria** | **Relevant Information from Institution Document (number of institutions)** |
| --- | --- |
| **Peer Reviewed Publications** | At least 3 publications. (n=1)  Continued scholarly publication (40 publications typically). (n=1)  Three articles for a 4 year contract; five articles for a 6-year contract. (n=1)  Three first authored papers published on important journals. (n=1)  One to three publications are required. (n=1)  Two publications per year are expected. (n=2)  Three to two papers required (varies based on impact factor of journals published in). (n=1)  Three to six papers required (varies based on impact factor of journals published in). (n=1)  Five to ten papers required (varies based on impact factor of journals published in). (n=2)  Two to four papers required (varies based on impact factor of journals published in, awards won, and citations). (n=1)  Two published papers required. (n=3)  Four published papers required. (n=1)  Five published papers since the last promotion. (n=2)  Six published papers required. (n=1)  Seven published papers required. (n=3)  Eight published papers required. (n=3)  Ten published papers required. (n=1)  Twelve published papers required. (n=1)  Twenty-eight publications preferred. (n=1)  One to six papers required (varies based on impact factor of journals published in). (n=1)  Requirement varies based on field of medicine, ranging from 14 - 53 publications in the last 10 years. (n=3) |
| **Authorship Order** | First or senior author on 1/3 of manuscripts. (n=1)  Greater points earned for single author, first author, or corresponding author on papers. (n=1)  Sole or lead authored publications are encouraged. (n=1)  Three first authored papers published in important journals. (n=1)  Expectation that candidates for promotion will have at least two publications per year with a significant portion as first or last author. (n=1)  Substantial research activity [..] as evidence by a substantial list of first or senior authored high quality peer-reviewed publications. (n=3)  Two papers as first or corresponding author. (n=1)  Six papers as first or corresponding author. (n=2)  Seven papers as first or corresponding author. (n=1)  Eight papers as first or corresponding author. (n=1)  Publishes as primary, senior, or corresponding author at least twice per year. (n=1)  At least 1 paper as first author. (n=1)  At least 1 sole authored paper. (n=1)  Five papers as first or corresponding author. (n=1)  First author on one of five papers. (n=1)  The specific role, such as lead author or otherwise, should be noted. (n=1)  Candidate will be expected to be lead, senior or corresponding author on most recent publications. (n=1)  Lead contributor or corresponding contributor of publications. (n=1)  Promotion requires sole, major, or senior authorship of high-impact publications. (n=1)  Corresponding author of at least five publications since previous promotion. (n=1)  Recent publications which the applicant is first or senior author. (n=1)  Applicant must be leading author (first or last) in the last three years. (n=1)  Independence as documented by senior authorship. (n=1)  At least 5 accredited publications with some as first author. (n=1) |
| **Impact Factor** | At least one paper with impact factor ≥3.0; at least one paper with impact factor ≥5.0, or cumulate impact factor rates totalling to ≥11.0. (n=1)  At least one paper with impact factor ≥9.0; or cumulate impact factor rates totalling to ≥30.0. (n=1)  At least one paper with impact factor higher than the average level of the specific area (if too few publications as first or senior author the impact factor must be higher). (n=1)  At least three paper with impact factors ranked as second level or higher on the journal citation reports. (n=1)  At least three papers with impact factor ≥5.0; at least one paper with impact factor ≥10.0, or cumulate impact factor rates totalling to ≥20.0. (n=1)  At least one paper with impact factor ≥10.0; or at least one paper with impact factor ≥15.0. (n=1)  At least three papers after the last evaluation with a cumulative impact factor of ≥4.0. (n=1)  At least 60% of papers in journals that have an impact factor. (n=1)  At least one paper in a Journal Citation Reports II area; or two papers with impact factor ≥3.0. (n=1)  *Table provided from guideline with varying requirements for impact factor dependent on the journal.* (n=1)  At least 7 papers with total impact factor ≥5.0; one paper with an impact factor ≥3.0; or one paper in journal citation reports II area. A second option is that papers reach an impact factor of at least 2 on the journal citation reports. A third option is that the professor has at least 7 relevant papers as first or corresponding author with all of the impact factors ≥3.0. (n=1) |
| **Grant Funding** | Principal investigator of one provincial (or higher level) grant. (n=1)  Five to seven years of funding. (n=1)  Principal investigator of one provincial (or higher level) grant; principal investigator of one national science foundation general project; principal investigator of vertical projects with total amount reach 500,000 RMB or above; or principal investigator of one national science foundation youth project; or principal investigator of vertical projects with total amount reach 300,000 RMB or above. (n=1)  Acquiring a multi-year extramural grant award as a principal investigator at the funding level of NIH R01, or higher; acquiring as a principal investigator a grant award comparable to NIH R01 from other agencies or foundations. (n=1)  Two national grants or one with 1,800,000 RMB. (n=1)  Principal investigator of one national science foundation general project; or principal investigator of sub-project of National Science and Technology Major Project; or principal investigator of National Key Research and Development Program, with research fund 1,000,000 RMB; or 500,000 RMB for non-provincial key disciplines. (n=1)  Principal investigator of two national natural science foundation project; or Principal investigator of project with more than 2,000,000 RMB.  Principal investigator of one national project and some publications. (n=1)  At least 1 grant funded since the last promotion.  Awarded with national second prize or provincial first prize; or expenses for the technology transfers reached 100,000 RMB, or scientific achievement application created more than 500,000 RMB, or gained 1000,000 RMB economic benefit. These could replace the requirements for the publication. (n=1)  Principal investigator of category A project with disposable funding amounting 250,000 for level four professors, 400,000 for level three professors, and 600,000 for level two professors. (n=1)  Principal investigator of one National Natural Science Foundation project. (n=1)  Must be principal investigator of two national grants. (n=1) |
| **Citations** | Threshold required for number of citations in the last 10 years for promotion. H-index threshold for the past 10 years also required, ranges from 5-12. (n=1)  Threshold required for number of citations in the last 10 years ranging from 16 - 406 based on medical specialty. H-index threshold for the past 10 years also required, ranges from 5-12. (n=2)  One paper need to be cited 20 to 30 times. Or 100- 150 cumulative citations among less than five papers. (n=1)  […] published one SCI paper with more than 30 times citation by others. (n=1)  "Citations from 10 hetero citations." (n=1) |

***Note*:** Data from non-English schools has been translated English.
