## Supplementary material for "Academic criteria for promotion and tenure in faculties of biomedical sciences: a cross-sectional analysis of 146 universities": Figure 1

**Figure 1. Included Universities**

**Available Documents**

Random sample of institutions (n= 170)

Guidelines not available (n= 60)

- Institution did not respond after two attempts to contact (n= 43)
- Institution stated that no guidelines are used for promotion and tenure (n= 4)
- Institution provided hiring documents only (n= 8)
- Institution not willing to share promotion and tenure guidelines (n= 2)
- Guidelines only available for positions that were not reviewed in the current study (n= 3)

Guidelines available (n= 110)

Guidelines available (n= 92)

- Faculty guideline (n= 39)
- Institution or national guideline (n= 53)

No faculty of biomedical sciences (n= 18)
