## Supplementary material for "Academic criteria for promotion and tenure in faculties of biomedical sciences: a cross-sectional analysis of 146 universities": Figure 2

**Figure 2. Mean Number of Criteria Present by Level of Promotion**

Traditional and Progressive Incentives for Promotion and Tenure

Median Number of Criteria

| Level of Evaluation | Traditional Incentives | Progressive Incentives |
| --- | --- | --- |
| Assistant Professor | 2 | 1 |
| Associate Professor | 2 | 1 |
| Professor | 3 | 1 |
| Tenure | 2 | 1 |
