## Supplementary material for "Academic criteria for promotion and tenure in faculties of biomedical sciences: a cross-sectional analysis of 146 universities": Table 1

**Table 1.** **Summary Characteristics of Included Institutions**

| **Variables** | **Institutions with Guidelines Available (n=92)** | **Institutions with Guidelines not Available (n=54)** | ***p value*** |
| --- | --- | --- | --- |
| **Criteria Level Available**  Institution (%)  Faculty (%) | 53 (58)  39 (42) | N/A | N/A |
| **Mean Leiden Ranking** **(SD)** | 358 (233) | 419 (251) | .327 |
| **Continent**  South America (%)  Australia (%)  Europe (%)  North America (%)  Asia (%)  Africa (%) | 1 (17)  6 (100)  27 (50)  28 (97)  29 (58)  1 (100) | 5 (83)  0 (0)  27 (50)  1 (3)  21 (42)  0 (0) | **.001** |
| **Human Development Index**  Very High (%)  High (%)  Medium (%) | 68 (64)  23 (62)  1 (50) | 39 (36)  14 (38)  1 (50) | .918 |
