## Supplementary material for "Academic criteria for promotion and tenure in faculties of biomedical sciences: a cross-sectional analysis of 146 universities": Table 2

| Table 2. Criteria of Interest for Promotion and Tenure | | | | | | | | |
| --- | --- | --- | --- | --- | --- | --- | --- | --- |
| Criteria | Presence of Criteria for Assistant Professor, n(%) | Presence of Criteria for Associate Professor, n(%) | Presence of Criteria for Full Professor, n(%) | Presence of Criteria for Tenure, n(%) | | Example of Quantitative Information if Present | | Example of Relevant Quote From University Website |
| Traditional Incentives |  |  |  |  | |  | |  |
| 1. Is any quantitative or qualitative mention made about publications required? If quantitative, please specify the requirement. | 39 (80%) | 76 (96%) | 79 (95%) | 22 (85%) | | “Minimum 2 research papers or one paper and one book” | | "Publication in refereed journals or series, or by publishers recognized as leaders in the field" |
| 2. Is any quantitative or qualitative mention made about the specific authorship order in publications? If so, please specify order (e.g. first, senior, single) required. | 11 (22%) | 28 (35%) | 28 (34%) | 9 (35%) | | “Three papers as first or corresponding author” | | “In case of multi-authored work, at least one of the peer reviewed publications must be sole authored." |
| 3. Is any mention made of journal impact factors? If quantitative, what are the minimum thresholds? | 12 (25%) | 24 (30%) | 23 (28%) | 2 (8%) | | “At least one impact factor>3.0; at least one impact factor>5.0, or accumulate impact factor>11.0” | | "At least 3 publications in international journals with reasonable impact factor are required." |
| 4. Is any mention made of grant funding? If quantitative, what are the minimum thresholds (i.e., amount of funding and/or number of grants as principal investigator)? | 26 (53%) | 50 (63%) | 56 (67%) | 15 (58%) | | “Principal investigator of one provincial project, principal investigator or the main member (top three) of project with more than 500,000 RMB” | | "The candidate has engaged in research grants/contracts as Principal or Co-Investigator at a funding level appropriate to the discipline, possibly in collaboration with other Universities or organizations" |
| 5. Is any mention made requiring that research is recognized at a national or international level? If so, please specify the requirement. | 11 (22%) | 26 (33%) | 39 (47%) | 11 (42%) | | N/A | | “Performance of exceptional distinction and achievements that are recognized as distinguished internationally or nationally (meeting the benchmarks)" |
| Progressive Evidence-Based Incentives |  |  |  |  | |  | |  |
| 6. Is any mention made of citations? If quantitative, what are the thresholds of minimum requirement? Are specific citation databases mentioned? | 12 (24%) | 23 (29%) | 23 (28%) | 6 (23%) | | “One paper cited more than 20 times” | | "Achieves a citation rate or proportion of research outputs in most prestigious outlets […] in line with discipline and leading universities" |
| 7. Is any mention made of data sharing? If quantitative, what are the minimum thresholds (e.g., percentage of data that is to be made available)? | 1 (2%) | 1 (1%) | 1 (1%) | 0 (0%) | | N/A | | "Sound data management is a basic requirement for this (academic analysis) and provides additional guarantees for a flawless methodology, for sharing and reusing data by other researchers in an Open Science context and for the accountability of a researchers own academic integrity" |
| 8. Is any mention made of publishing in open access mediums? If quantitative, what are the minimum thresholds (e.g., percentage of studies to be published in open access journals)? | 0 (0%) | 0 (0%) | 0 (0%) | 0 (0%) | | N/A | | N/A |
| 9. Is any mention made of registration (including preregistration challenge) of studies? If yes, are there thresholds of minimum requirement (e.g., percentage of studies that are to be registered). | 0 (0%) | 0 (0%) | 0 (0%) | 0 (0%) | | N/A | | N/A |
| 10. Is any mention made of adherence to reporting guidelines for publications? If so, are specific guidelines mentioned? | 0 (0%) | 0 (0%) | 0 (0%) | 0 (0%) | | N/A | | N/A |
| 11. Is any mention made of alternative metrics for sharing research (e.g., social media and print media)? If so, are specific metrics mentioned? | 3 (6%) | 3 (4%) | 2 (2%) | 1 (4%) | | N/A | | "Ghent University has invested in a proper information system of research output (biblio, IWETO/FRIS) for many years and is currently extended this to other research-related areas (Gismo). As soon as publications and activities are properly registered in the information system, the administrative burden on researchers in the context of an evaluation (promotion, project applications) should be reduced to a minimum." |
| 12. Is any mention made of accommodations or adjustments to expectations due to extenuating circumstance? If so, please specify the description of accommodations (e.g., an extra year to defer tenure consideration) and the type of eligible circumstances (e.g., parental leave, medical leave)? | 22 (45%) | 28 (35%) | 29 (35%) | | 13 (50%) | "A female faculty […] may extend her contract up to two years in association with pregnancy and delivery and up to one year in case of adopting a child six years old or younger." | "Where staff have had a career break, long term absence or other extenuating circumstances which impact on their output/performance, they are encouraged to provide this information including what impact such breaks/absences have had on them undertaking their role." | |
