## Supplementary material for "Academic criteria for promotion and tenure in faculties of biomedical sciences: a cross-sectional analysis of 146 universities": Table 3

**Table 3. Multiple Linear Regression Analyses of Institution Characteristics and Presence of Traditional and Progressive Criteria** (N = 83).

| **Variables** | **Traditional Criteria** | | | **Progressive Criteria** | | |
| --- | --- | --- | --- | --- | --- | --- |
|  | **Unstandardized β**  **(95% CI)** | **t** | **p-value** | **Unstandardized β**  **(95% CI)** | **t** | **p-value** |
| **Criteria Level** (Reference group = Faculty Level) | -.60 (1.45 to 3.22) | -1.89 | .062 | -.08 (-.48 to .31) | -.42 | .675 |
| **Leiden Ranking** | .01 (-.001 to .001) | -.15 | .884 | .01 (-.001 to .001) | -.45 | .656 |
| **Human Development Index (HDI)** (Reference group = Very High HDI)  Medium HDI |  |  |  |  |  |  |
|  | -- | -- | -- | -- | -- | -- |
| High HDI | .63 (-.19 to 1.45) | 1.52 | .132 | -.29 (-.81 to .22) | -1.14 | .260 |
| **Continent** (Reference group = Asia)  South America  Australia  Europe  North America  Africa |  |  |  |  |  |  |
|  | -1.29 (-3.64 to 1.07) | -1.09 | .279 | -.45 (-1.92 to 1.01) | -.62 | .540 |
|  | 1.78 (.58 to 2.99) | 2.94 | .**004** | 1.07 (.32 to 1.82) | 2.85 | .006 |
|  | .48 (-.41 to 1.36) | 1.08 | .286 | .32 (-.23 to .86) | 1.15 | .256 |
|  | .99 (.13 to 1.84) | 2.29 | .025 | .43 (-.11 to .96) | 1.59 | .115 |
|  | 1.70 (-.64 to 4.05) | 1.45 | .151 | -.47 (-1.93 to .99) | -.64 | .526 |

*Note:* bolded confidence intervals = p < .005
