## Supplementary material for "Academic criteria for promotion and tenure in faculties of biomedical sciences: a cross-sectional analysis of 146 universities": Table 4

**Table 4. Logistic Regression Analyses of Institution Characteristics and Presence of Traditional and Progressive Criteria** (N = 83).

| **Variables** | | **Criteria Present** | **Criteria Not Present** |  | | |
| --- | --- | --- | --- | --- | --- | --- |
|  | | **N (%)** | **N (%)** | **Odds Ratio** | **95% Confidence Intervals** | |
|  | | | | | **Lower** | **Upper** |
| **Mention of authorship order** | | 28 (33.7) | 55 (66.3) |  |  |  |
|  | **Criteria level** (Reference group = faculty level) | 17 (60.7) | 19 (34.5) | .35 | .10 | 1.22 |
|  | Institution Level | 11 (39.3) | 36 (65.6) |  |  |  |
|  | **Leiden Ranking** [Mean (SD)] | 336.1 (290.2) | 371.5 (214.6) | 1.00 | 1.00 | 1.00 |
|  | **Human Development Index** (HDI) **(**Reference group = Very High HDI)  Medium HDI  High HDI | 18 (64.3) | 43 (78.2) |  |  |  |
|  |  | 0 (0.0) | 1 (1.8) | .00 | .00 | -- |
|  |  | 10 (35.7) | 11 (20.0) | 1.53 | .30 | 7.69 |
|  | **Continent** (Reference group =Asia)  South America  Australia  Europe  North America  Africa | 12 (42.9) | 14 (25.5) |  |  |  |
|  |  | 0 (0.0) | 1 (1.8) | .00 | .00 | -- |
|  |  | 2 (7.1) | 4 (7.3) | 1.21 | .12 | 12.61 |
|  |  | 3 (10.7) | 19 (34.5) | .33 | .05 | 2.22 |
|  |  | 11 (39.3) | 16 (29.1) | .73 | .14 | 3.87 |
|  |  | 0 (0.0) | 1 (1.8) | .00 | .00 | -- |
| **Mention of journal impact factor** | | 23 (27.7) | 60 (72.3) |  |  |  |
|  | **Criteria level** (Reference group = faculty level) | 10 (43.5) | 26 (43.3) | .73 | .18 | 2.86 |
|  | Institution Level | 13 (56.5) | 34 (56.7) |  |  |  |
|  | **Leiden Ranking** | 327.8 (252.0) | 371.8 (238.5) | 1.00 | 1.00 | 1.00 |
|  | **Human Development Index** (HDI) **(**Reference group = Very High HDI)  Medium HDI | 11 (47.8) | 50 (83.3) |  |  |  |
|  |  | 1 (4.3) | 0 (0.0) | 4.45 | .00 | -- |
|  | High HDI | 11 (47.8) | 10 (16.7) | 3.35 | .65 | 17.41 |
|  | **Continent** (Reference group = Asia)  South America  Australia  Europe  North America  Africa | 12 (52.2) | 14 (23.3) |  |  |  |
|  |  | 0 (0.0) | 1 (1.7) | .00 | .00 | -- |
|  |  | 2 (8.7) | 4 (6.7) | 1.50 | .13 | 16.87 |
|  |  | 5 (21.7) | 17 (28.3) | .84 | .13 | 5.27 |
|  |  | 3 (13.0) | 24 (40.0) | .30 | .04 | 2.03 |
|  |  | 1 (4.3) | 0 (0.0) | .00 | .00 | -- |
| **Mention of grant funding** | | 56 (67.5) | 27 (32.5) |  |  |  |
|  | **Criteria level** (Reference group = faculty level) | 28 (50.0) | 8 (29.6) | .49 | .13 | 1.84 |
|  | Institution Level | 28 (50.0) | 19 (70.4) |  |  |  |
|  | **Leiden Ranking** | 354.0 (260.6) | 371.0 (200.4) | 1.00 | 1.00 | 1.00 |
|  | **Human Development Index** (HDI) **(**Reference group = Very High HDI)  Medium HDI  High HDI | 43 (76.8) | 18 (66.7) |  |  |  |
|  |  | 1 (1.8) | 0 (0.0) | 1.63 | .00 | -- |
|  |  | 12 (21.4) | 9 (33.3) | 1.89 | .38 | 9.49 |
|  | **Continent** (Reference group = Asia)  South America  Australia  Europe  North America  Africa | 13 (23.2) | 13 (48.1) |  |  |  |
|  |  | 0 (0.0) | 1 (3.7) | .00 | .00 | -- |
|  |  | 6 (10.7) | 0 (0.0) | 3.61 | .00 | -- |
|  |  | 12 (21.4) | 10 (37.0) | 2.40 | .44 | 13.10 |
|  |  | 24 (42.9) | 3 (11.1) | 9.96 | 1.52 | 65.31 |
|  |  | 1 (1.8) | 0 (0.0) | .00 | .00 | -- |
| **Mention of national or international recognition** | | 39 (47.0) | 44 (53.0) |  |  |  |
| **Criteria level** (Reference group = faculty level) | | 21 (53.8) | 15 (34.1) | .49 | .11 | 2.26 |
| Institution Level | | 18 (46.2) | 29 (65.9) |  |  |  |
| **Leiden Ranking** | | 356.3 (250.5) | 362.4 (236.2) | 1.00 | 1.00 | 1.00 |
| **Human Development Index** (HDI) **(**Reference group = Very High HDI)  Medium HDI  High HDI | | 36 (92.3) | 25 (56.8) |  |  |  |
|  |  | 1 (2.6) | 0 (0.0) | 7.68 | .00 | -- |
|  |  | 2 (5.1) | 19 (43.2) | .47 | .05 | 4.14 |
| **Continent** (reference group =Asia)  South America  Australia  Europe  North America  Africa | | 2 (5.1) | 24 (54.5) |  |  |  |
|  |  | 0 (0.0) | 1 (2.3) | .00 | .00 | -- |
|  |  | 5 (12.8) | 1 (2.3) | 52.36 | 2.30 | 1190.95 |
|  |  | 9 (23.1) | 13 (29.5) | 6.86 | .71 | 66.85 |
|  |  | 22 (56.4) | 5 (11.4) | 27.44 | **3.26** | **231.16** |
|  |  | 1 (2.6) | 0 (0.0) | .00 | .00 | -- |
| **Mention of citations** | | 23 (27.7) | 60 (72.3) |  |  |  |
| **Criteria level** (Reference group = faculty level) | | 8 (34.8) | 28 (46.7) | .65 | .16 | 2.75 |
| Institution Level | | 15 (65.2) | 32 (53.3) |  |  |  |
| **Leiden Ranking** | | 316.7 (257.8) | 376.0 (235.1) | 1.00 | 1.00 | 1.00 |
| **Human Development Index** (HDI) **(**Reference group = Very High HDI)  Medium HDI  High HDI | | 19 (82.6) | 42 (70.0) |  |  |  |
|  |  | 0 (0.0) | 1 (1.7) | .00 | .00 | -- |
|  |  | 4 (17.4) | 17 (28.3) | 1.64 | .22 | 12.53 |
| **Continent** (reference group =Asia)  South America  Australia  Europe  North America | |  |  |  |  |  |
|  |  | 4 (17.4) | 22 (36.7) |  |  |  |
|  |  | 0 (0.0) | 1 (1.7) | .00 | .00 | -- |
|  |  | 5 (21.7) | 1 (1.7) | 47.81 | 2.25 | 1016.33 |
|  |  | 8 (34.8) | 14 (23.3) | 5.21 | .58 | 46.80 |
|  |  | 6 (26.1) | 21 (35.0) | 1.97 | .24 | 16.33 |
| Africa | | 0 (0.0) | 1 (1.7) | .00 | .00 | -- |
| **Mention of accommodations/adjustments to circumstances** | | 29 (34.9) | 54 (65.1) |  |  |  |
| **Criteria level** (Reference group = faculty level) | | 15 (51.7) | 21 (38.9) | .93 | .19 | 4.45 |
| Institution Level | | 14 (48.3) | 33 (61.1) |  |  |  |
| **Leiden Ranking** | | 353.5 (258.6) | 362.8 (234.3) | 1.00 | 1.00 | 1.00 |
| **Human Development Index** (HDI) **(**Reference group = Very High HDI)  Medium HDI  High HDI | | 29 (100.0) | 32 (59.3) |  |  |  |
|  |  | 0 (0.0) | 1 (1.9) | .00 | .00 | -- |
|  |  | 0 (0.0) | 21 (38.9) | .00 | .00 | -- |
| **Continent** (reference group =Asia)  South America  Australia  Europe  North America  Africa | | 2 (6.9) | 24 (44.4) |  |  |  |
|  |  | 0 (0.0) | 1 (1.9) | .00 | .00 | -- |
|  |  | 4 (13.8) | 2 (3.7) | 5.07 | .45 | 57.79 |
|  |  | 6 (20.7) | 16 (29.6) | 1.01 | .15 | 6.95 |
|  |  | 17 (58.6) | 10 (18.5) | 4.54 | .62 | 33.44 |
|  |  | 0 (0.0) | 1 (1.9) | .00 | .00 | -- |

*Note:* bolded confidence intervals = p < .005
